# Global Trends in eHealth Research: Analysis and Visualization of Author and Indexer-Supplied Keywords

**DOI:** 10.1101/2020.11.26.399881

**Authors:** Williams Ezinwa Nwagwu, Omwoye Bosire Onyancha

## Abstract

This article examined the growth and development of ehealth research based on the headcount and analysis of the characteristics of keywords used by authors and indexers to represent their research content during 1945-2019 as indexed in the Elsevier’s Scopus database. The results show that although the term ehealth originated in the late 1990s, but it has become an envelope term for much older terms such as telemedicine, and its variants which originated much earlier. The keywords were spread in 27 Scopus Subject Areas, with medicine (44.04%), engineering (12.84%) and computer science (11.47%) leading while by Scopus All Science Journal Classification, Health Sciences accounted for 55.83% of the keywords and physical sciences followed with 30.62%. The rest two classifications namely social sciences and life sciences made only single digit contributions. Although the primary essence of ehealth was how to meet health needs, the work of engineers who either initially deployed telephone to meet their health needs or, and, computer scientists, who addressed the need to design technologies for medical services is very significant. It is concluded that ehealth is a multidisciplinary area that is attractive to researchers from all disciplines because of its sensitive focus on health, and therefore requires pooling and integrating of resources and expertise, methods and approaches.

## 1.0 Introduction

Timely and efficient transfer, and or, exchange of information can make a tremendous difference in various health situations - whether the situation is an emergency due to an accident, say, or a lifelong chronic health condition, or other (Thomas 2018, WHO 2018, Dorsey and Tople 2020). Advances in information and communication technology (ICT) have facilitated successful sharing of data thereby enhancing universal access to information for health products and services. Globally, the health sector has been implementing a variety of ICTs to improve the efficiency of information exchange at all healthcare levels. Besides exchanging of information, modern ICTs have also facilitated clinical and consultation services of medical practitioners for timely and cost effective healthcare delivery. This development is generally known as ehealth, that is, the use of ICT to meet for healthcare purposes (Liu, Su and Ji 2018). The applications in ehealth include interactive websites, e-mail, wearable technologies, telehealth applications, gaming, web portals, voice recognition, and online communities (Kreps and Neuhauser, 2010).

Research on ehealth is expanding due to its benefits. According to Mettler and Raptis (2012), e-health is directed to achieving quality, cost effective, equal and customized healthcare. E-health plays significant roles in, and contributes to health promotion, disease prevention and treatment (Wicks, Stamford, Grootenhuis, Haverman and Ahmed 2014, Chaomei 2017) and reduction of service costs and improvement of quality of health service. Minichiello, Rahman, Dune, Scott and Dowsett (2013) discussed increased user and supplier control of health intervention, and influence on government policy making as an advantage of ehealth. Generally, ehealth is making healthcare more efficient and allowing patients and professionals to access and manage data in ways that were previously impossible.

A crucial but yet scientifically inexhaustively explored aspect of scientific literature is the nature of accumulation of literature and its role in understanding the growth, and development of disciplines, and further implication for science and technology and human resources development. Since the introduction of the concept of ehealth in the literature in 1999 (Mitchell 1999), there has been extensive use and application of the concept. Ehealth concept should be evaluated to understand its clarity, and to know whether it is well-defined and differentiated from other concepts, and whether the literature definitions are consistent (Morse, Hupcey and Cerdas, 1996). The concept is fast maturing, with dedicated journals (for instance *JMIR*), conferences (Eysenbach 1999) and research institutes and centres, for instance Norwegian Centre for E-health Research (Skrøvseth and Laukli 2019), and also exists as a cognate discipline in many health institutions, departments in universities and government ministries. Spearheaded by WHO, there are collaborations within and across nations and regions geared towards facilitating learning, pooling of resources and fast tracking information exchange for efficient ehealth. WHO’s “eHealth unit works with partners at the global, regional and country level to promote and strengthen the use of ICT in health development, from applications in the field to global governance.” (WHO 2018e).

Bibliometrics/scientometrics provides us with the mathematical and statistical applications and methods to understand the quality and quantity, evolution, growth and development, and other aspects, of published scientific literature is necessary to understand the nature and structure of the subject of ehealth (Sweileh 2017). One way to examine the growth and development of a new discipline or concepts is to start from understanding the structure of the discipline or concept. The structure of a discipline or a concept includes the theoretical definition of the discipline, its attributes, boundaries, preconditions, and outcomes (Fridahl 2010). Examining the structure of a discipline requires, among others, understanding authors’ and indexers’ behaviours, for example, the keywords they choose to represent the content they create. To what extent do author supplied keywords locate ehealth as a medical, engineering, computer science, or other, subject? What do the keywords that represent ehealth documents tell us about the extent of reach or spread of the discipline? Addressing these questions is important for assessing ehealth maturity as a discipline, curriculum design and course content development (Onyancha 2019), and collaboration and disciplinary location. Basically, ehealth and the researches that have come on the subject in the past suggest a multidimensional combination of health with information technology towards achieving efficiency in the health system.

A keyword is a word or phrase that succinctly describes the contents of a particular document. It is the shortest generalized summary of a document and it serves as an important index for research papers. Author- and indexer-created keywords represent the opinion of the researcher regarding how best to represent a research paper using the shortest summary. They can also help researchers spot and locate a new area of knowledge. In fact keyword frequency could be accepted as an indication of consensus of researchers regarding the content and subject matter of a new or even old study area. Keywords serve to enhance the representation of contents. The principal significance of keywords is that they serve as a tool for retrieving literature; they are guideposts to the subject the researcher considers to be in focus in a study. Examining the growth of author supplied keywords provides an innovative way of examining a biological property of literature, namely growth and development of the literature often reflected and influenced by environmental factors, and subject matter. A census of keywords could provide useful information about the nature and characteristics of knowledge in a subject area, and provide further basis for possible prediction of future attention. Such a headcount in could also provide an insight into the dependence of research interest over time – time being strongly linked to social, economic and political circumstances of the embrace of a subject. Blessinger and Frasier (2010) have demonstrated that the evidence base of the role and significance of keywords in scientific literature is well developed in library and information science.

This paper examined the maturity of e-health concept in the literature based on keywords used by authors to represent their research, covering the maximum period during which ehealth has existed in global literature (1945-2019). A mature concept is a concept that is “well defined, has clearly described characteristics, delineated boundaries, and documented preconditions and outcomes” (Morse, et al., 1996, p. 387). The paper explored the ehealth literature development from the perspective of clusters, links, link strength and frequency of the author and indexer selected keywords, using information visualization technology to understand growth and evolution of the subject. Various approaches were adopted to classify the keywords, and subject areas and classes of the documents in order to unveil latent information about the subject matter. Progressively, the study synthesized links among the keywords, and constructed and visualized analytics of the structural dynamics, trends and patterns to understand the formation and evolution of ehealth hot topics, and the trend of the subject growth.

How do the keywords used by the authors represent their evaluation of the content of their papers? How does the structure of the keywords reflect a body of intellectually corrigible subject matter consisting of cognate concepts, facts and theories? What does the cluster of the keywords tell us about ehealth – are scholars approaching a convergence in respect of the new discipline? Do the links suggest restricted discipline with high degree of linkage between different research areas within the discipline or unrestricted discipline with relatively diffuse links within and outside the discipline? Does the community of scholar engaging on ehealth research coalesce around any central intellectual and content agenda? What further development would be required to refine the subject such that scholars in the area represent a research community that has internal communication paraphernalia, for example universities, and professional societies? The point of emphasis though is not the superiority or necessity of disciplinarity, and the need therefore for ehealth to become a granular area of knowledge that is disconnected from other disciplines. The point rather is to establish the structure of ehealth in order to understand what is new, what is borrowed and what exact social problems ehealth solves.

### Objectives of the study

This study was designed to:

1. map the global pattern of growth of research on ehealth during 1945-2019 using author- and indexer supplied keywords of papers published in the area,
2. examine most innovative keyword words and the survival rate of the keyword, and,
3. analyse the broad subjects scope of the area in order to determine disciplinary structure and characteristics, and disciplinary sources of influence,

## 2.0 Literature Review

To properly situate this study, we undertook the literature review from three perspectives (i) clarification of concepts (ii) review of some related empirical studies and (iii) review of theoretical perspectives focusing on evolution of disciplines.

### Concept Clarification and some Timeline

Some concept clarification and timeline is necessary in this study where there are many concepts that are interwoven in their meanings and operations, but the concepts at the same time have different histories. Concpet clarification is necessary to properly interpret the keywords and the subjects they represent as they guide us to understand the growth, maturity and boundaries of ehealth. The key concepts that require some clarification are ehealth, telemedicine and mobile health; their variants will be assumed or mentioned where necessary.

#### The Concept of eHealth

The concept of ehealth actually emerged in the 1990s as one of the e-terms that the explosion of the internet brought about (Mitchell 1999). With the email and other electronic support technologies happening during this time, the possibilities of multidimensional communication compelled the attachment of the *e* to almost every existing term: email, ecommerce, ehealth etc. The term ehealth could be said to be well understood to the extent that it refers to health; but this understanding is from a broad perspective as its precise meaning, disciplinary independence and spread of constituency are still evolving. Many researchers, industries, organizations and The World Health Organisation have since engaged the concept (WHO 2016, 2018, Eysenbach 2001, Liu et al 2019). In 2012, WHO defined “eHealth as the cost-effective and secure use of information and communication technologies (ICTs) for health and health-related fields”, and in 2018 as “the use of information and communication technologies (ICT) for health” (WHO 2018, p.1). Moving from a relatively more specific definition to a very broad, flexible and elastic one must be for the purpose of accommodating roles for a wide array of both present and future technologies, including those in existence before the arrival of the concept for efficient addressing of health needs.

In its current definitions, ehealth encompasses the range of uses of information technologies from primarily health records purposes to those roles that are often considered best suited for face to face encounters such as diagnosis, drug prescription, and examination, among others. Liu, Su and Ji (2018) summarized it this way: “With the development of information technology, e-health has been absorbing and applying emerging information technologies and applications” (p.8). Besides emerging information technologies, the concept has also absorbed older technologies that existed before the birth of the internet and the WWW.Generally therefore, ehealth is a broad concept used to describe all electronic applications, telehealth and mobile health, and all their variants for the purpose of meeting health needs.

#### Concept of Telemedicine and Telehealth

Telemedicine and telehealth are often used interchangeably although there exists a slight difference. According to Vladzymyrskyy (2016) “Telemedicine encompasses diagnostic, treatment and prevention processes within the frame of modern health care services, which are carried out primarily by means of telecommunication and computer technologies” (p.1). Thomas (2018) makes it shorter, “Telemedicine refers to the provision of remote clinical services via real-time two-way communication between the patient and the healthcare provider through electronic audio and visual means” (p.1). The World Health Organisation has earlier defined telemedicine as “The delivery of health care services, where distance is a critical factor, by all health care professionals using information and communication technologies for the exchange of valid information for diagnosis, treatment and prevention of disease and injuries, research and evaluation, and for the continuing education of health care providers, all in the interests of advancing the health of individuals and their communities” (WHO, 2010 p.1).

On its own part, telehealth is the delivery and facilitation of health and health-related services including medical care, provider and patient education, health information services, and self-care via telecommunications and digital communication technologies. NEJM Catalyst (2019) puts the relationship this way: “…telemedicine refers specifically to the practice of medicine via remote means, telehealth is a blanket term that covers all components and activities of healthcare and the healthcare system that are conducted through telecommunications technology.” (p1).

Telemedicine is, in practice, older than the telephone. Vladzymyrskyy (2016) has revealed that telemedicine was in existence long before the invention of telephone. He described a telemedical device called sphygmosphone created by Dr Jabez Baxter Upham and his colleagues in 1858. Sphygmosphone was used to fix the heart pulse as a curve and then the data was sent as a telegraph. By 1859, the device was tested and successfully used to transmit the heart rate data of a patient suffering congenital sternal fissure from one hospital to another. The invention of the telephone boosted telemedicine. According to Aronson (1977) “A principal and ongoing research since the invention of the telephone by Graham Bell and his colleagues in 1876 has been how to bridge the gap between patients and healthcare givers. In fact, this challenge became one of the major uses to which the telephone was put when it was finally invented” (p.12)” The *Lancet* published a large number of articles focusing on the telephone and healthcare during 1876 to 1975; telephone was first mentioned in Lancet in the issue of 9 February 1878. In a letter to the editors, an author had suggested that the telephone could improve medical diagnosis and that it could be useful in demonstrating and studying the sound produced by a muscle during contact, the negative contraction, among others. In this regard, the idea of using the telephone for clinical and consultation services such as auscultation, and home management of non-critical emergencies, and the advantages they portend were initiated early in the history of *ehealth*.

#### The Concept of Mobile Health

m-Health has been defined as “… medical and public health practice supported by mobile devices, such as mobile phones, patient monitoring devices, personal digital assistants (PDAs), and other wireless devices” (Adibi 2012 p1). In another resource, WHO (2016) defined mhealth exactly the same way it defined ehealth, as “…use of mobile wireless technologies for public health,” capturing the role of wider range of mobile technologies. The critical significance of mobile health is that mobile wireless technologies enable individuals to carry along with them access to healthcare services they need. Mobile health/medicine is often conflated with telemedicine/telehealth (Thomas 2018) because mobile health is tied to telecom technologies.

### Some Empirical Bibliometric Studies on eHealth Keywords

As has been suggested earlier, ehealth encompasses all electronic technology applications in health. There exists ample evidence of efforts to charactertise ehealth literature. Fatehi and Wootton (2012) examined the occurrence of the terms ‘telemedicine’, ‘telehealth’ and ‘e-health’ in the Scopus database. They found a total of 11,644 documents containing one of the three terms in the title or abstract. They also found that telemedicine was the most common term, with 8028 documents referring to it, followed by e-health (2573) and then telehealth (1679). According to them, documents with telemedicine in their titles or abstracts first appeared in 1972, and have continued to appear at a low rate until 1994 when they started to increase rapidly. They also observed that the growth of telehealth only began to increase about five years later. In his own study Groneberg (2015) carried out a scientometric and density equalizing analysis of telemedicine based on data collected from WoS. They found that during the period from 1900 to 2006 a number of 3290 items were identified, and that the first publication was in telemedicine was in 1964. They concluded that in all subject categories examined for published items related to telemedicine, *healthcare sciences and services* ranked first by far, followed by *medical informatics* and *medicine, general and internal*.

González et al (2018) performed an author keyword analysis for mapping Sport Sciences using data mining technique. They conducted an analysis of the frequency of appearance and the dynamics of the author keywords and constructed a network of co-occurrences of the keywords and the survival time of new words that have appeared since 2001 has also been analysed. They concluded that sports science is increasingly multidisciplinary and that the word *rehabilitation* sort of colonizes the field. Gupta, Dhawan and Mueen (2018) undertook a scientometric assessment of global publications on digital health research output during 2007–2016. The study covered 6981 publications sourced from Scopus database. They found that medicine is the most studied subject with largest publication share in digital health research, followed by computer science, engineering, health profession, and others. In Fang’s (2015) scientometric review of the structure and the evolving of digital medicine, he defined digital medicine as “… an interdiscipline which integrated computer science, information engineering with medicine, digital medicine originally and mainly on digital medical imaging technology research for accuracy and speedy clinical diagnosis and therapy” (p.25). According to them, the earliest cluster was on medical imaging segmentation and registration, and then post-processing imaging technology, detector, phase contrast, reversible watermarking, input, model 3D reconstruct, real-time dynamic imaging, dosimetry have been the hot research topics. They also said that the internet health information is a recent cluster is on.

Liu, Si and Ji (2019) conducted a bibliometric study to detect and characterize ehealth research during 2001–2016. Based on keywords used in the 6371 documents in their study, they classed the categories and ranked the subjects into (i) Internet technology; (ii) telemedicine, telehealth, m-health, and communication (iii) randomized control trial, (iv) healthcare field and (v) health management. They also identified research directions on ehealth: healthcare science and service, computer science, medical informatics, engineering, public environmental occupational health, telecommunications, psychology, general internal medicine, and information science library science. They found that the articles involve some elements of clinical areas such as nursing, cancer treatment, pharmacy, and science and technology development. Based on their study, they suggested that the top four research directions on ehealth are healthcare science and services, computer science, medical informatics and engineering.

Yang (2019) carried out a diachronic keyword analysis in research article titles and cited article titles in applied linguistics from 1990 to 2016. They collected and investigated titles written in leading applied linguistics journals over 25 years to identify their keywords. They compared their data over different time periods to study the significance of the domain knowledge. They found that keywords varied according to the research trends and that titles of articles are getting longer as more keywords are deployed by authors in order to increase the visibility of the paper and enhance the citability of the papers. Onyancha (2019) examined the evolution of information literacy over 43 years (from 1975 to 2018), and visualized and mapped knowledge of its literature in the Scopus database based on keyword analysis. He found that information literacy has evolved from being a library- and/or librarianship-oriented concept to a multidisciplinary field that is no longer restricted to social sciences but spreads across 27 disciplines in Scopus’ subject classification.

Ahmadvand *et al* (2019) have conducted a bibliographic-bibliometric analysis on articles published in *Journal of Medical Internet Research* that used “digital health” as a keyword and evaluated the trends, topics, and citations of the publications during January 2000 and August 2019. They found 1797 articles having “digital health” as a keyword, and they were mostly published between 2016 and 2019. Of these articles 277 articles (32.3%) were published by *Journal of Medical Internet Research* and the most frequently used keyword for was *mhealth*.

By way of synthesis, (i) telemedicine came into the health and medical literature before 1876 when telephone was invented, long before the e-revolution of the 1990s, and it was first used for medical/consulting purpose. Its modern day use has been traced by Vladzymyrskyy (2016) to the work of Gale (1927) in a new article, and in a scientific literature by Murphy (1970). (ii) mobile health came into the literature in 1945 during the first world war and referred to mobility of physical health infrastructure including human to meet human health challenges; its modern day version which refers to how information technologies could bridge the gap between the patient and the healthcare giver only came into existence in 1974 (iii) the concept *ehealth* was born in the 1990s but the expansion of ICTs has positioned ehealth as a clearing house concept for all information technology based healthcare services, products and systems. Without any doubt therefore all the tele-and mobile, and other health technologies and their variants are components of ehealth (Sweileh (2017).

## 3.0 Research methodology

The source of the data for the current study was the Elsevier’s Scopus database. The database is one of the largest bibliographic databases in the world besides the Clarivate Analytics’ Web of Science products. The database has become a key source of bibliographic and citation data for bibliometric and scientometric studies. The following search query involving ehealth and its variations was conducted to obtain relevant data.

((KEY (ehealth) OR KEY ({e-health}) OR KEY ({e health}) OR KEY ({electronic health}) OR KEY (telehealth) OR KEY ({tele-health}))) OR ((KEY (emedicine) OR KEY ({e-medicine}) OR KEY ({e medicine}) OR KEY (“electronic medic*”) OR KEY (telemedicine) OR KEY ({tele-medicine}) OR KEY ({mobile medicine}) OR KEY ({m-medicine}))) OR ((KEY (mhealth) OR KEY ({m-health}) OR KEY ({m health}) OR KEY ({mobile health}))) AND (EXCLUDE (PUBYEAR, 2020)) AND (LIMIT-TO (DOCTYPE, “ar”) OR LIMIT-TO (DOCTYPE, “cp”))

Multiple search terms were used because researchers’ choice of keywords vary very significantly. For instance, researchers may use the keyword smartphone, texting, cellular phone or mobile application and join them with any word in the field of health to enable them more specifically represent the content of their research. The search for ehealth research publications was limited to conference papers and research articles published before 2020. *Errata* documents and corrections of published articles were excluded from the analysis because they would not represent actual publications. Also, only conference papers that appeared under source type were retained because they would not appear again as published papers, thus avoiding duplication in publications.

The search yielded a total of 86186 documents and 82968 keywords. The data was exported to MS Excel in .csv format and saved for analysis. Publications on ehealth were available for 62 years during the 75 years of the coverage of the study (1945-2019), indicating somewhat constant and consistent interest of researchers on the subject matter, albeit truncated or segmented during certain years. Table 1 shows the volume of keywords per annum.

**Table 1:** Number of keywords per year.

| Year | No | % | Year | No | % |
| --- | --- | --- | --- | --- | --- |
| 1945 | 1 | 0,00 | 1989 | 62 | 0,07 |
| 1946 | 1 | 0,00 | 1990 | 90 | 0,11 |
| 1947 | 2 | 0,00 | 1991 | 68 | 0,08 |
| 1949 | 2 | 0,00 | 1992 | 116 | 0,14 |
| 1950 | 1 | 0,00 | 1993 | 160 | 0,19 |
| 1960 | 2 | 0,00 | 1994 | 245 | 0,29 |
| 1962 | 1 | 0,00 | 1995 | 445 | 0,53 |
| 1965 | 2 | 0,00 | 1996 | 429 | 0,52 |
| 1966 | 2 | 0,00 | 1997 | 570 | 0,69 |
| 1967 | 2 | 0,00 | 1998 | 716 | 0,86 |
| 1968 | 5 | 0,01 | 1999 | 691 | 0,83 |
| 1969 | 24 | 0,03 | 2000 | 739 | 0,89 |
| 1970 | 74 | 0,09 | 2001 | 680 | 0,82 |
| 1971 | 74 | 0,09 | 2002 | 739 | 0,89 |
| 1972 | 106 | 0,13 | 2003 | 901 | 1,08 |
| 1973 | 84 | 0,10 | 2004 | 976 | 1,17 |
| 1974 | 79 | 0,09 | 2005 | 1248 | 1,50 |
| 1975 | 97 | 0,12 | 2006 | 1575 | 1,89 |
| 1976 | 82 | 0,10 | 2007 | 1851 | 2,23 |
| 1977 | 48 | 0,06 | 2008 | 2114 | 2,54 |
| 1978 | 72 | 0,09 | 2009 | 2663 | 3,20 |
| 1979 | 62 | 0,07 | 2010 | 3148 | 3,78 |
| 1980 | 55 | 0,07 | 2011 | 3963 | 4,76 |
| 1981 | 33 | 0,04 | 2012 | 4613 | 5,55 |
| 1982 | 38 | 0,05 | 2013 | 5358 | 6,44 |
| 1983 | 37 | 0,04 | 2014 | 6154 | 7,40 |
| 1984 | 38 | 0,05 | 2015 | 6854 | 8,24 |
| 1985 | 61 | 0,07 | 2016 | 7126 | 8,57 |
| 1986 | 46 | 0,06 | 2017 | 8158 | 9,81 |
| 1987 | 57 | 0,07 | 2018 | 9022 | 10,85 |
| 1988 | 61 | 0,07 | 2019 | 10462 | 12,58 |

The data was grouped and analyzed according to four time-periods, i.e. 1945-1990, 1991-2000, 2001-2010 and 2011-2019 so as to assess the development and evolution of ehealth research during the period (see table 2). We obtained the broad subject categories within which the ehealth documents are indexed using the *analyze search results* function provided on the Scopus results platform so as to determine the fields that have contributed the most to ehealth research.

**Table 2:** Number of keywords per document per period

| Periods | Total number of keywords | Total number of documents | Keywords per document |
| --- | --- | --- | --- |
| 1945-1990 | 159 | 1402 | 0.11 |
| 1991-2000 | 2653 | 4179 | 0.63 |
| 2001-2010 | 16724 | 15895 | 1.05 |
| 2011-2019 | 65432 | 61710 | 1.06 |
| Total | 82968 | 86186 | 0.96 |

VOSviewer software was used to analyse the data according to the author- and indexer-supplied keywords and map the development and evolution of ehealth research over time. VOSviewer is widely used to perform different types of analyses including co-authorship, co-occurrence, citation analysis, bibliographic coupling, and co-citation analysis. The software allows one to perform co-occurrence analysis using three units of analysis namely all keywords, author keywords or index keywords. The author- and indexer supplied keywords, which constituted the unit of analysis for the current study, have been extensively used in bibliometric studies (Onyancha 2019). Graphical visualization methods are effective in discovering network patterns. Information visualization offers one a quick and independent, scientific judgment of the objective evidence of data (Tho, Yeung, Wei, Chan and So 2017, Xiao, Li, Sun and Zhang (2017). However, maps have inadequate capability in discovering spatial and temporal patterns of connections in a network especially when the network exists and changes across space and time (Koylu et al 2014).

## Results

The study covers a period of 75 years (1945-2019) but data was only available for 62 years with entries for the periods 1946 and 1951 to 1959, and, 1961 and 1963-1964. Keywords were consistently supplied for papers published from 1967 up and until 2019. With particular respect to the author- and indexer-supplied keywords, Table 1 demonstrates that the number of keywords has continued to grow, from just 0.11 keywords per paper in 1945-1990 to 1.03 in 2011-2019. Overall, the average number of author keywords per paper for ehealth research is below one.

### The formative period: 1945-1990

There was a total of 1402 documents on ehealth during this period; but only 44 of this large volume of documents supplied keywords, a total of 143 or 0.19 keywords per document and about 6 keywords per year. The significance of keywords in enhancing accessibility and usability of documents was not yet strongly realised as at this period, and there also appears not to be any rules demanding that authors represent their research with keywords. Towards the end of this period however, the electronic mail, the WWW and the internet had been established as potent information technologies that can drive healthcare, and use of keywords in research has matured significantly.

The term “mobile health” came into the literature in 1960 when Cachia (1960) published “A Mobile Health Unit amongst the Masai” in *East African Medical Journal,* although *mobile health* did not appear in the author keywords. The concept of mobile health at that time was not used in its present day connotation “…as the application of mobile phones and other small, portable and wireless computing and communication devices to meet the information and service needs of healthcare providers and clients” (WHO 2017 p.2). Rather, the focus was on how to take healthcare facilities to meet people’s health needs in their own locations, and not how information technologies could leverage healthcare access between providers and patients.

**Figure 1.** shows that no keyword is dominant among the 143 keywords that captured the contents of the documents during the period, but *public health/hospitals* and *clinics* were somewhat outstanding. Table 3 shows the ten keywords that appeared in two or more documents, and illustrates the pattern of uptake of ehealth research during this period. Table 1 shows further that the numbers of keywords per document from 1945 to 1967 lie between zero and two for each year, and 5 in 1968. Keywords started growing from 1969, with no truncation to the peak date of 2019. For the period under consideration, there was a gentle growth from 1969, a spike in 1972, and a continuous moderate but steady growth from 1973. There was a total of 143 keywords for 1402 documents in the 1945-1990 time period. Out of these, 10 appeared in the literature twice or more times. Of particular interest is the occurrence of telemedicine and mobile health services during the 1945-1990 time period. The other keywords namely *breast cancer, community dentistry, emergency medical services, epidemiology, mass screening, mobile unit and public health/hospitals* and *clinics* depict the focus areas in which telemedicine and mobile health services were applied.

Furthermore, of the ten keywords (see table 3), *public health/hospitals and clinics* has the highest number of clusters but zero links and zero link strength. Public health is a very wide area of health encompassing education, policy making and research, disease and injury prevention using surveillance, prevention, preparedness, and health promotion strategies. *Emergency medical services* has only one cluster but it has the highest number of links and also the highest total link strengths (27 apiece). It can be observed that the keywords that emerged as prominent during 1945-1990 excluded the keywords of the documents that heralded the birth of ehealth, for instance hospitals/military, hospitals/mobile, and surgery/operating rooms. It is worth repeating that during this period, the use of the word “mobile” and its appendages described movement of health facilities and resources to the location of the sick.

**Table 3:** Author and indexer supplied keywords 1945-1990

| No. | Label | Cluster | Links | Total link strength | Frequency |
| --- | --- | --- | --- | --- | --- |
| 1 | Cardiac arrest | 4 | 8 | 8 | 3 |
| 2 | Breast cancer | 2 | 6 | 6 | 2 |
| 3 | Community dentistry | 7 | 5 | 5 | 2 |
| 4 | emergency medical services | 1 | 27 | 27 | 2 |
| 5 | Epidemiology | 12 | 4 | 4 | 2 |
| 6 | Mass screening | 2 | 6 | 7 | 2 |
| 7 | Mobile | 2 | 4 | 4 | 2 |
| 8 | Mobile unit | 2 | 6 | 7 | 2 |
| 9 | Public health/hospitals and clinics | 32 | 0 | 0 | 2 |
| 10 | Telemedicine | 5 | 6 | 6 | 2 |

Besides the study of Jacob (1965) which introduced the term *medical records systems*, the seminal work of Holzer (1974) on telemedicine announced the arrival of research and possibilities in the role of *tele-* technologies in healthcare. He defined telemedicine as “… as the practice of medicine at long distance by means of telecommunications, in particular, closed-circuit television and telemetry” (p5). Jacob’s research focussed on a “Two-way television enables physicians to establish a nominal doctor-patient relationship with patients at a remote location, while providing the means for visual examination of patients” (p2). Mayo-Wells (1963) has rendered telemetry as “….the collection of measurements or other data at remote or inaccessible points and their automatic transmission to receiving equipment for monitoring” (p1). The word *telemetry* is not very common in the health literature today, probably because of rapid development of micro and more diverse telecommunication and information technology facilities, for instance, GSM, and others which have provided more efficient ways of collecting and sharing data among patients, and, healthcare providers. From the study of Segal (2015) however it could be inferred that abnormal heart activities are a major aspect of healthcare where telemetry has been mostly applied. It is not surprising therefore that *cardiac arrest* has the highest frequency in the top ten keywords during 1945-1990.

### The Period of Growth and Development: 1991-2000

The period 1991-2000 was consciously tagged the period of growth and development where growth is used in its normative sense of linear numerical increase in various aspects of the concept. This period also marked some remarkable development on the role of information technologies in healthcare, particularly the emergence of mobile technologies, and rapid expansion in information technologies; but the term *electronic health* is still absent from the author keywords used to represent studies during the period. However, the following e-health related keywords appeared in the documents published in this period: electronic health records, electronic healthcare records, electronic medical records, electronic medication monitoring, mobile healthcare, mobile health clinics, and mobile health units, thereby signaling the presence of ehealth in the period despite the absence of the actual term.

**Figure 2.** shows that compared to the previous period, the visualization map is more densely clustered, with shorter and thicker links, signifying more attention to the subject and more intensity of research, based on the deployment of keywords, during the period. There is also emerging significance of research on *telemedicine* because it has the largest label and the highest weight, signifying the critical importance of the keyword in ehealth research.

Ehealth research during the period generated 4179 documents in ten years, accounting for 42 documents per year and 0.63 keywords per document. While this result points to increase in research in the area, there is need to observe that the practice of deploying keywords to represent key content of research appears to be more embraced during this period than the previous.

Table 4 displays the top 30 keywords that had at least a frequency of 9, used to represent content of research papers in this period. *Telemedicine* is the only term in cluster one that appeared in nine and above documents, with a frequency of 399, 100 links and 545 total link strength and as such illustrates how much research has focused on the envisaged role of telephones in healthcare. Telemedicine services have replaced phone calls from general practitioners to specialists for advice, and travel for many patients. The opinions of Allen (2000) is informative: Allen (2000) “… telemedicine remains linked to medical professionals, while e-health is driven by non-professionals, namely patients (or, in the e-health jargon, consumers) that with their interests drive new services even in the healthcare field-mostly for their empowerment through access to information and knowledge” (p12). Nesbith (2014) reported *teleradiology* as an ancillary telemedicine service, but the two clusters of the technology has a frequency of 69, a total link strength of 168 and was connected to 53 other items. The single cluster of *internet* whose third rank in the list of keywords could be considered significant because the frequency of 59 is considerably high for a technology that was only escalated in the middle of the 80’s, and the total link strength of 110 and 39 links signify the emergence of a technology that would fast-track applications of electronic technologies for healthcare. Except education, evaluation, quality assurance, security and ISDN, much of the keywords were directly related to application of telecommunication and information technology to health.

**Table 4:** Major Keywords in ehealth during 1991-2000.

| No. | Label | Cluster | Links | Total link strength | Frequency |
| --- | --- | --- | --- | --- | --- |
| 1 | Telemedicine | 1 | 100 | 545 | 399 |
| 2 | Teleradiology | 2 | 53 | 168 | 69 |
| 3 | Internet | 1 | 39 | 110 | 56 |
| 4 | Telepathology | 1 | 21 | 84 | 41 |
| 5 | Electronic medical record | 3 | 14 | 20 | 35 |
| 6 | PACS | 2 | 31 | 85 | 29 |
| 7 | Multimedia | 4 | 23 | 42 | 21 |
| 8 | Dicom | 2 | 21 | 55 | 17 |
| 9 | ISDN | 1 | 17 | 44 | 17 |
| 10 | Telecommunications | 1 | 25 | 44 | 16 |
| 11 | Telehealth | 1 | 12 | 20 | 16 |
| 12 | Virtual reality | 1 | 15 | 24 | 16 |
| 13 | Evaluation | 2 | 15 | 32 | 14 |
| 14 | Medical imaging | 2 | 16 | 44 | 14 |
| 15 | Medical informatics | 3 | 16 | 29 | 14 |
| 16 | Security | 4 | 20 | 32 | 14 |
| 17 | Radiology | 2 | 20 | 43 | 12 |
| 18 | Teleconsultation | 1 | 7 | 15 | 12 |
| 19 | World Wide Web | 1 | 13 | 24 | 12 |
| 20 | Electronic medical records | 3 | 9 | 10 | 11 |
| 21 | Telematics | 1 | 14 | 23 | 11 |
| 22 | Computers | 3 | 16 | 23 | 10 |
| 23 | Education | 1 | 15 | 25 | 10 |
| 24 | Screening | 5 | 8 | 18 | 10 |
| 25 | Ultrasound | 2 | 11 | 18 | 10 |
| 26 | Information technology | 1 | 9 | 11 | 9 |
| 27 | Quality assurance | 3 | 10 | 14 | 9 |
| 28 | Remote consultation | 1 | 14 | 24 | 9 |
| 29 | Tele-medicine | 3 | 3 | 5 | 9 |
| 30 | Video conferencing | 1 | 13 | 23 | 9 |

### The Period of Expansion - 2001-2010

During this period, what is generally known as modern information technology and its applications have become very mature, and their applications in health have also become outstandingly mature. The peak of this expansion was in this period was marked by the successful implementation of video chats using Skype and other chat applications, a development that has tremendously boosted ehealth and ehealth research. Table 5 shows the 30 keywords that have a frequency of at least 58. Liu Su and Ji (2019) observed that in 2001– 2005, the link intensity among high-frequency keywords was low. They also observed that the study of ehealth was at an exploratory stage, and research direction was scattered because scholars had not yet formed a complete theoretical system. But the emergence of ehealth concepts has raised great academic interest and scholars were beginning to use network communication technology which greatly improved the quality of medical services and reduced healthcare costs (Eysenbach 2008).

**Figure 3.** shows that *Telemedicine* remains the largest label with the highest weight; and table 3 shows that it has a single cluster and occurred 1820 times, and generated 97 links and a large total link strength of 1379 which are demonstrative of dominant a keyword on ehealth research. We interpret this observation to mean that the teletechnologies is very outstanding in place of increasing convergence between computing and, telecommunications networks and media content in addition to other information technologies to healthcare. Liu, Si and Ji (2019) had the same result when they showed telemedicine was the dominant keyword that during 2001-2005, and 2006-2010, but that the trend changed in 2011-2016 when *internet* became the most dominant. Liu Su and Ji (2019) said:

**Table 5:** Keywords used in ehealth research during 2001-2010.

| No. | Label | Cluster | Links | Total link strength | Frequency |
| --- | --- | --- | --- | --- | --- |
| 1 | Telemedicine | 1 | 97 | 1379 | 1820 |
| 2 | E-health | 1 | 71 | 436 | 495 |
| 3 | Telehealth | 1 | 60 | 313 | 318 |
| 4 | Internet | 1 | 66 | 321 | 259 |
| 5 | Electronic health record | 2 | 52 | 209 | 255 |
| 6 | Ehealth | 2 | 58 | 214 | 225 |
| 7 | Electronic medical record | 2 | 55 | 160 | 204 |
| 8 | Electronic health records | 2 | 54 | 160 | 196 |
| 9 | Electronic medical records | 2 | 52 | 132 | 181 |
| 10 | Healthcare | 2 | 50 | 165 | 131 |
| 11 | Security | 2 | 39 | 175 | 117 |
| 12 | Medical informatics | 3 | 48 | 136 | 109 |
| 13 | Primary care | 2 | 42 | 108 | 108 |
| 14 | Privacy | 2 | 38 | 161 | 107 |
| 15 | Information technology | 2 | 43 | 143 | 92 |
| 16 | Interoperability | 2 | 33 | 117 | 85 |
| 17 | Telemonitoring | 1 | 36 | 100 | 84 |
| 18 | Diabetes | 1 | 41 | 93 | 76 |
| 19 | Evaluation | 3 | 34 | 92 | 73 |
| 20 | Technology | 1 | 40 | 107 | 72 |
| 21 | Information systems | 3 | 47 | 112 | 71 |
| 22 | Healthcare | 3 | 39 | 85 | 70 |
| 23 | ECG | 1 | 21 | 71 | 67 |
| 24 | Home care | 1 | 31 | 91 | 61 |
| 25 | Her | 2 | 31 | 77 | 60 |
| 26 | Teleradiology | 1 | 30 | 87 | 60 |
| 27 | Electronic patient record | 2 | 25 | 45 | 59 |
| 28 | Hypertension | 1 | 26 | 59 | 59 |
| 29 | Standards | 2 | 39 | 102 | 59 |
| 30 | Medical records | 3 | 33 | 67 | 58 |

*In 2006–2010, with the Internet explosively developing and governments attaching more importance to E-health gradually, some medical items based on network technology entered the implementation phase. Scholars tried to evaluate implementation of these projects from visual map aspects. The formation of E-health research prototype has an important connection with the Internet, telemedicine, and care (p8).*

Despite Mea’s (2001) pessimism in the paper in *JMIR* titled *What is eHealth: The Death of Telemedicine* in which he anticipated that the birth of ehealth would make telemedicine less weighty in describing computer and information technology-oriented healthcare, telemedicine has remained a strong concept during the period. According to Mea (2001), “…in the context of a broad availability of medical information systems that can interconnect and communicate - telemedicine will no longer exist as a specific field. The same could also be said for any other traditional field in medical informatics, including information systems and electronic patient records. e-Health presents itself as a common name for all such technological fields” (p12). Rather, components of telemedicine such as *teleradiology* and *telemonitoring* among others are occurring as keywords in ehealth research, underpinning the persisting significance of telemedicine.

One could pick out the keyword *private,* used in relation to a study addressing how ehealth could help individuals manage their private health issues; and so do other new keywords such as *diabetes* and *hypertension*, among others, illustrate various aspects of ehealth research application. The keyword *standards* arose from the point of view of a researcher advocating for standards with respect to ehealth applications.

The expansion in the concept of ehealth has been generally attributed to the role of *Journal of Internet Medical Research* (JMIR) of Eysenbach and team in 1999 (Al-Rimawi et al. 2016). The keyword *e-health* which was to become an umbrella word for concepts that existed before it, for instance, telemedicine, came into the literature for the first during this period, and has grown rapidly during the 2000s. With its label being nearly as large as telemedicine and a frequency of 495, the concept occurred in cluster one and has 71 links and a strong total link strength of 436 within about its first ten years of birth. It must also be observed the variant of the concept, *ehealth,* which appeared as a cognate keyword support the emerging compromise about the role of the keyword in describing technology oriented healthcare. It can be inferred from the work of Eng (2001) that the overwhelming *e* in *health* is attributable to the increasing role of the internet in technology convergence. Some authors, for instance, Pretlow (2007) and Car, Black, Anandan, Cresswell and Pagliari, (2009) have defined ehealth by highlighting and paying specific attention to the role of the internet, bypassing the significance of telecommunication.

### The Period of Maturity: 2011-2019

eHealth has evidently matured during this period. Table 6 shows the list of keywords that appeared at least 400 times as against 58 in the previous period. The ten year period generated a total of 61710 keywords or an average of 6171 keywords per year (see table 5). Despite evidence of increasing research in the area (Dorsey and Tople 2020), there is also observed increase in the inclusion of keywords for research papers. This contrasts with the former periods when keywords appeared not to be compulsory inclusions as representatives of content of research papers. Figure shows that although telemedicine remains weightier in terms of size of label, the weight has reduced in this period compared with the previous. According to Dorsey and Tople (2020) “The past decade saw telemedicine finally cross this chasm. In the USA, at least 15% of physicians work in practices that use telemedicine and adoption by private insurers increased by 50% per year for most of the decade” (p16). In a previous study that covered the period 2015 to 2017, Barnett, Ray and Souza (2018) have observed a very high volume of use of telemedicine services for telemental and primary healthcare purposes in the United States. Wootton and Bonnardot (2015) made the same observation in developing countries, despite Fatehi and Wootton’s (2012) study which showed that research in telemedicine is more intense in developed countries than in the developing countries.

**Figure 4.** shows that the weight of the label *telemedicine* has reduced drastically in comparison with the previous period. This is despite *telemedicine* having a frequency of 4828, occurring in three clusters, with 106 links and a total link strength of 4364, figures that are almost three times more the previous period (see table 5). How does one explain the reduction in the weight of the telemedicine label despite the high indices? Basically one can see that the statistics for all the keywords have very high indices, indicating that research on ehealth during this period is not only intense, but it is also heavily spread to many other knowledge areas.

**Table 6:** Keywords used in ehealth research during 2011-2019.

| No | Label | Cluster | Links | Total link strength | Frequency |
| --- | --- | --- | --- | --- | --- |
| 1 | Telemedicine | 3 | 106 | 4364 | 4828 |
| 2 | Electronic health records | 1 | 102 | 2391 | 2825 |
| 3 | Mhealth | 2 | 105 | 3308 | 2578 |
| 4 | E-health | 3 | 106 | 2207 | 2321 |
| 5 | Ehealth | 2 | 106 | 2741 | 2147 |
| 6 | Telehealth | 3 | 102 | 2199 | 1810 |
| 7 | Electronic health record | 1 | 100 | 1343 | 1534 |
| 8 | Mobile health | 2 | 100 | 1710 | 1456 |
| 9 | Electronic medical records | 1 | 94 | 685 | 849 |
| 10 | Primary care | 1 | 95 | 1048 | 795 |
| 11 | Electronic medical record | 1 | 92 | 620 | 767 |
| 12 | Healthcare | 4 | 87 | 711 | 584 |
| 13 | Diabetes | 2 | 99 | 867 | 581 |
| 14 | Smartphone | 2 | 89 | 831 | 552 |
| 15 | Machine learning | 1 | 81 | 627 | 548 |
| 16 | Internet | 2 | 86 | 906 | 538 |
| 17 | Technology | 3 | 94 | 904 | 536 |
| 18 | M-health | 3 | 83 | 576 | 516 |
| 19 | Depression | 2 | 82 | 757 | 504 |
| 20 | Privacy | 4 | 57 | 673 | 484 |
| 21 | Natural language processing | 1 | 64 | 456 | 460 |
| 22 | Health information technology | 1 | 81 | 658 | 456 |
| 23 | Self-management | 2 | 78 | 896 | 449 |
| 24 | Stroke | 3 | 76 | 462 | 432 |
| 25 | Quality improvement | 1 | 77 | 443 | 428 |
| 26 | Security | 4 | 46 | 599 | 422 |
| 27 | Primary healthcare | 1 | 75 | 546 | 411 |
| 28 | Patient safety | 1 | 67 | 443 | 410 |
| 29 | Epidemiology | 1 | 65 | 342 | 404 |
| 30 | Medical informatics | 1 | 73 | 506 | 400 |

Opinions of researchers that define ehealth mainly from electronic medical records perspective (Pearce and Haikerwal 2010) became weighty in the study as the keyword *electronic health records* and its variants increased in their use. Also, *mhealth* and its variants were absent in the list of top 30 keywords in the previous period, but they have appeared as important keywords during this current period.

## Analysis by Scopus Broad Subject Areas

Scopus has classified disciplines into 32 subject areas. Based on table 7, 44% of the keywords belonged to the subject of medicine while 13% and 11 % respectively belonged to the subjects of computer science and engineering while 32% of the keywords are distributed to the rest of the 27 subject areas (minus undefined). This result immediately shows the pattern of the emerging subject - while researchers in medicine are concerned with how to use existing information technologies at that time to further the practice of the diagnosis and treatment, engineers were busy constructing tools and computer scientists were testing models. Generally, Pharmacology, Toxicology and Pharmaceutics; Psychology, and, Decision Sciences as well as Business Management and Accounting, agricultural and biological sciences and earth and planetary sciences are latecomers to research on ehealth. The multidisciplinary area did not also supply any keywords until the third period of the study. The undefined category

**Table 7:** Scopus Subject Classes.

|  |  | **1945-1990** | **1991-2000** | **2001-2010** | **2011-2019** | **N** | **%** |
| --- | --- | --- | --- | --- | --- | --- | --- |
| 1 | Medicine | 745 | 3477 | 11192 | 44043 | 59457 | 44.04 |
| 2 | Computer Science | 218 | 191 | 3536 | 13403 | 17348 | 12.85 |
| 3 | Engineering | 268 | 1110 | 3877 | 10227 | 15482 | 11.47 |
| 4 | Health Professions | 61 | 324 | 2549 | 7188 | 10122 | 7.50 |
| 5 | Nursing | 42 | 188 | 1164 | 3812 | 5206 | 3.86 |
| 6 | Biochemistry, Genetics and Molecular Biology | 57 | 159 | 603 | 3745 | 4564 | 3.38 |
| 7 | Social Sciences | 13 | 28 | 640 | 2737 | 3418 | 2.53 |
| 8 | Mathematics | 48 | 112 | 485 | 2085 | 2730 | 2.02 |
| 9 | Pharmacology, Toxicology and Pharmaceutics | 2 | 6 | 325 | 1653 | 1986 | 1.47 |
| 10 | Physics and Astronomy | 61 | 133 | 335 | 1305 | 1834 | 1.36 |
| 11 | Psychology | 1 | 5 | 186 | 1398 | 1590 | 1.18 |
| 12 | Neuroscience | 19 | 12 | 153 | 1185 | 1369 | 1.01 |
| 13 | Materials Science | 68 | 179 | 316 | 798 | 1361 | 1.00 |
| 14 | Decision Sciences | 2 | 12 | 194 | 1041 | 1249 | 0.93 |
| 15 | Chemical Engineering | 18 | 69 | 372 | 684 | 1143 | 0.85 |
| 16 | Business, Management and Accounting | 1 | 16 | 207 | 757 | 981 | 0.73 |
| 17 | Agricultural and Biological Sciences | 1 | 1 | 31 | 857 | 890 | 0.66 |
| 18 | Immunology and Microbiology | 2 | 22 | 66 | 685 | 775 | 0.57 |
| 19 | Multidisciplinary | 0 | 0 | 31 | 745 | 776 | 0.57 |
| 20 | Arts and Humanities | 14 | 11 | 95 | 442 | 562 | 0.42 |
| 21 | Environmental Science | 9 | 21 | 22 | 406 | 458 | 0.34 |
| 22 | Energy | 3 | 4 | 64 | 356 | 427 | 0.32 |
| 23 | Chemistry | 15 | 11 | 40 | 354 | 420 | 0.31 |
| 24 | Dentistry | 12 | 82 | 65 | 214 | 373 | 0.28 |
| 25 | Veterinary | 1 | 4 | 27 | 167 | 199 | 0.15 |
| 26 | Earth and Planetary Sciences | 0 | 7 | 49 | 74 | 130 | 0.10 |
| 27 | Economics, Econometrics and Finance | 3 | 1 | 33 | 82 | 119 | 0.08 |
| 28 | Undefined | 19 | 3 | 0 | 9 | 31 | 0.02 |
| 29 | **Total** | **1402** | **6188** | **26657** | **42841** | **77389** | 100 |
| 30 | **%** | 2.20 | 7.10 | 34.45 | 55.36 |  |  |

During the early period of ehealth research (1945-1991), the concept had already spread across 25 different subject areas (minus the undefined category). Vellsen et al (2013) has described ehealth as a multidisciplinary subject while in his integrative review of ehealth interventions, Jannsen et al (2017) referred to ehealth as interdisciplinary. Besides supporting that different disciplines are teaming up to address the problem of health through technology, table 6 shows that ehealth has become an attractive intellectual tourist site that provides a research space for researchers from various subjects. All the same, ehealth is domestic in *medicine* which accounted for 745 out of the 1702 documents, followed by engineering (268) and computer science (218) during 1945-1990.

Medicine also dominates the subject classes during 1991-2000 absorbing 3477 keywords while engineering and health professions follow with 1110 and 324 keywords respectively and computer science remains within the first four top subject areas (191), out of the total of 6188 keywords generated during the period. During 2001-2010, of the 26657 documents generated, the volume of keywords that belong to the subject of medicine continued to increase accounting for 11192 or 42% of the total, engineering accounted for 3877 or 14% while computer science took the third place (3536) keywords or 13%. Finally, during 2011-2019, medicine accounted for 44043 keywords or 44% of the total 100452 keywords, and computer science rising in share of keywords (13403 or 13%) while engineering accounted for 10227 or 10 of the keywords. Generally, of the 135000 documents, medicine accounted for about 77% with computer science and engineering absorbing 22% and 20% respectively (figure 6).

## Analysis by Scopus All Science Journal Classification (ASJCC)

Scopus has classified subjects into four main classes known as All Science Journal Classification Codes namely physical sciences, health sciences, social sciences and life sciences. The essence of this section is to understand the subject classes that the keywords used by ehealth researchers actually represent; The Scopus All Science Journal Classification enables us see clearer how each subject area performed within a classification. This presentation enables us reflect on the nature of contributions to ehealth when the consideration is with regards to Health Science, Physical Sciences, Social Sciences and Life Sciences, each of these classes has been defined by Scopus.

Table 8 is very revealing about attention to ehealth. Figure 7 shows that more than half (55.83%) of the documents belong to the Health Sciences class while 30.62% belong to the Physical Sciences. Life Sciences accounted for 7.09% while Social Sciences accounted only for 5.87%.

**Table 8:** Scopus All Science Journal Classification.

| Subject Area | Subject Area Classifications | 1945-1990 | 1991-2000 | 2001-2010 | 2011-2019 | Total (%) | Overall Total % |
| --- | --- | --- | --- | --- | --- | --- | --- |
| Physical Sciences | Chemical Engineering | 18 | 69 | 372 | 684 | 1143(0.85) | 30.62 |
|  | Chemistry | 15 | 11 | 40 | 354 | 420(0.31) |  |
|  | Computer Science | 218 | 191 | 3536 | 13403 | 17348(12.85) |  |
|  | Earth and Planetary Sciences | 0 | 7 | 49 | 74 | 130(0.10) |  |
|  | Energy | 3 | 4 | 64 | 356 | 427(0.32) |  |
|  | Engineering | 268 | 1110 | 3877 | 10227 | 15482(11.47) |  |
|  | Environmental Science | 9 | 21 | 22 | 406 | 458(0.34) |  |
|  | Material Science | 68 | 179 | 316 | 798 | 1361(1.00) |  |
|  | Mathematics | 48 | 112 | 485 | 2085 | 2730(2.02) |  |
|  | Physics and Astronomy | 61 | 133 | 335 | 1305 | 1834(1.36) |  |
|  | Multidisciplinary | - | - | - | - | -- |  |
| Health Sciences | Medicine | 745 | 3477 | 11192 | 44043 | 59457(44.04) | 55.83 |
|  | Nursing | 42 | 188 | 1164 | 3812 | 5206(3.86) |  |
|  | Veterinary | 1 | 4 | 27 | 167 | 199(0.15) |  |
|  | Dentistry | 12 | 82 | 65 | 214 | 373(0.28) |  |
|  | Health Professions | 61 | 324 | 2549 | 7188 | 10122(7.50) |  |
|  | Multidisciplinary | - | - | - | - | -- |  |
| Social Sciences | Arts and Humanities | 14 | 11 | 95 | 442 | 562(0.42) | 5.87 |
|  | Business, Management and Accounting | 1 | 16 | 207 | 757 | 981(0.73) |  |
|  | Decision Sciences | 2 | 12 | 194 | 1041 | 1249(0.93) |  |
|  | Economics, Econometrics and Finance | 3 | 1 | 33 | 82 | 119(0.08) |  |
|  | Psychology | 1 | 5 | 186 | 1398 | 1590(1.18) |  |
|  | Social Sciences | 13 | 28 | 640 | 2737 | 3418(2.53) |  |
|  | Multidisciplinary | - | - | - | - | -- |  |
| Life Sciences | Agricultural and Biological Sciences | 1 | 1 | 31 | 857 | 890(0.66) | 7.09 |
|  | Biochemistry, Genetics and Molecular Biology | 57 | 159 | 603 | 3745 | 4564(3.38) |  |
|  | Immunology and Microbiology | 2 | 22 | 66 | 685 | 775(0.57) |  |
|  | Neuroscience | 19 | 12 | 153 | 1185 | 1369(1.01) |  |
|  | Pharmacology, Toxicology and Pharmaceutics | 2 | 6 | 325 | 1653 | 1986(1.47) |  |
|  | Multidisciplinary | - | - | - | - | - |  |

In the Physical Sciences Class, understandably computer science (12.85%) and engineering (11.47%) led in contribution. While engineers are engaged in design of technologies, computer scientists are concerned with models for solving problems. Earth and Planetary Sciences were late comers in the area and they made the least contribution (0.10%), Energy and Environmental Sciences accounted for only 0.3%. Medicine and Health professions remain leaders in their Class accounting for 44% and 7.5% respectively while Veterinary (0.15) has the least contribution to the Class. Social Sciences is the subject area with the highest volume of contribution in the Social Sciences Class (2.53%) while Biochemistry, Genetics and Molecular Biology (3.53%) has the highest contribution from the Life Sciences Class.

## Conclusive Remarks

This study was designed to examine the maturity of ehealth through analysis of keywords in papers published in journals and conference proceedings during 1945-1990. The term *ehealth* contains two key themes namely *electronic* and *health;* all other terms used to describe this term are either descriptive of the themes, or are a further description of the concerns of an author in using the terms. Between the two themes *electronic* and *health* however, health is obviously the superior element. Otherwise why is it electronic that is reduced to *e*, and not health to *h*? Besides ehealth happening alongside the *e*revolution, “No doubt, throughout human evolution, health and diseases always were matters of main concern and had a profound effect on human society, shaping it” (Vladzymyrskyy 2016 p.iii).

Between and around *electronic* and *health* are a world of life, physical, social and health sciences realities that invite the attention of researchers from various subjects areas and classes, health being everybody’s social capital and wealth. In the couple *electronic health*, electronic is the driver of health. When *electronic* is used to drive *health,* what is the product? The deliverable is healthcare or health service in the form of multifaceted information such as transmission of glucometer, or stethoscope reading or other from the patient to the healthcare provider for consultation and counselling purposes. It could also be a piece of simple and direct information such as drug taking reminder or body temperature reading. *Electronic* in *ehealth* is all encompassing, an envelope of any modern technologies means adopted for healthcare; it could be the computer, digital, internet, social media, telephone, mobile technology, and chat etc. Primarily, ehealth involves technology, people and the health of the people within a technology and social system that operate in a milieu whose activities are organised and managed by government and non-government agencies, private and non-private organisations, corporate and other institutions, healthcare managers and various educational institutions. It involves the health of individuals and their health at work, home and community

As is also the case in this study, health is one of the most coupled themes; for instance, electronic health, public health, occupational health, and environmental health, etc. Most strikingly, it is only in the case of electronic health that the interpretation of the couple appears to depart from the way others are interpreted. For instance, public health can abstemiously be interpreted as the aspect of health that is concerned health of the public. We can perform such a permutation with other couples of health, except electronic health. eHealth is concerned with how healthcare professionals deliver their care and service, how patients or clients of healthcare providers access or procure or choose to, healthcare services; it is concerned with how the business people exploit the opportunity available in the technology and systems to make health services available and accessible to care givers and clients (Kovac, 2014, Cunningham et al., 2014). eHealth concerns itself with how the government, and the communities deploy or support the deployment of the couple of electronic health through healthcare experts and providers and organisations to meet their health target. eHealth concerns education and research, administration, health monitoring and surveillance and business and access to healthcare (Curran and Curran, 2005; Maheu et al., 2002). Besides clinical practices, promotional and prevention purposes, the focus of ehealth to specific diseases such as diabetes, and heart conditions, among others signals the potentials of ehealth in managing lifelong diseases. The essence of ehealth is human health.

The wide array of keywords used to describe research on ehealth shows that e-health pervades all aspects of, and invites the attention of all to, the healthcare delivery challenges (Kovac, 2014). They include all health disciplines and concerns whether consultation, promotion, rehabilitation, preventive, or others. They also include referrals, health management, and remote healthcare services at local, national or international levels; advocacy systems, collaborative networks and health administration. Ehealth also includes the concerns of psychologists and associated disciplines, commerce and businesses as well as engineers and computers science. Despite this wide array of interests, ehealth is primarily concerned with to deploy and use to meet the health people’s health challenges.

Although the term *ehealth* developed much later than telemedicine (and its variants), telemedicine remains the hottest topic on the subject. Most strikingly, the question of remote clinical services via real-time two-way communication between the patient and the healthcare provider through electronic audio and visual means was also the concern of earliest experiments.

## Implications of the study for Practice, Policy and Society

A bibliometric analysis and mapping of the keywords that represented *ehealth* literature 1945-2019 tell us so much about *ehealth* literature, and is significant for further development of the area. First, the effort to deploy information technologies for improved healthcare is older than many disciplines in medical field, as well as the new information technologies. Yet, *ehealth* is a new area of knowledge, with no clear conceptual standing and disciplinary location. Evidently the lack of convergence on the conceptual interpretation of *ehealth* poses a serious challenge to healthcare policy and education, including librarians whose job it to classify disciplines and provide access to information resources. eHealth is obviously predominantly a health science discipline given the huge attention it has received from the subjects areas, compared with the others. What it means is that despite the interests of researchers from various fields, ehealth should be located within the health science, with an unrestricted participation by researchers from other disciplines. The implication of this observation is the need for a multidisciplinary curriculum to facilitate efficient research and education, and, teaching and learning.

There is also need for collaborative strategies for non-research and use related issues and activities to organize, and ensure that, use of ehealth products and services. Early in the experiments of telemedicine, doctors have flagged what could be regarded as the possibility of telemedicine leading to abuse in medical processes, and unethical practices. For instance, is there a danger for patients preferring telephone consultation to an office at just the cost of the use of the technology? Aronson has cited a discussion on this as far back as 1883: “The only fear we have is that when people can open up a conversation with us for a penny, they will be apt to abuse the privilege, and that to have a dozen telephone consultations in one day, or conversations that might be thought to supersede a consultation, would be a doubtful addition to one’s advantage or repose.” (Cited in Aronson 1977 p.4). May (2011) and Ja *et al* (2009) have alluded to possible professional abuse that may have drastic consequences both for patients and the society. eHealth systems should be holistic, taking into consideration the business of the medical practitioner, psychological factors as well as ethical issues often involved in medical practice.

Shiferaw and Mehari (2020) have discussed the concept of ehealth literacy, an aspect of the discourse where collaborators should involve all stakeholders including library and information science. eHealth literacy has been conceptualized and defined as the the ability of internet users to locate, evaluate, and act upon web-based health information (<u>Tubaishat</u> and <u>Habiballah</u> 2016). Given the reservations shared by medical practitioners regarding the possible infraction of ehealth into professional practice, ehealth literacy should define very clearly what healthcare practices should be at the discretion of patients, and when to seek for a face to face consultation.

Paige et al (2018) have introduced a transaction model for ehealth literacy based on the observation that existing models have limited theoretical underpinnings that reflect the transactional capabilities and they classed ehealth literacy skillset as consisting functional, communicative, critical and translational components. This and other existing models are concerned with how patients will benefit maximally from ehealth, efforts on how ehealth should be practiced to avoid abuse of professional practice. Research on ehealth will continue, as the core of the concept is health and the focus of the researchers will continue to be diverse. The curial challenge remains how to bring together the outcomes from different researches and generate a multidisciplinary theoretical perspective that can guide further development. Research on ehealth will continue, as the core of the concept is health and the focus of the researchers will continue to be diverse. The key challenge remains how to bring together the outcomes from different researches and generate a multidisciplinary theoretical perspective that can guide further development.

In the course of writing up this paper it was discovered that there were studies on ehealth that used the term *digital health, cybermedicine* and many of the *tele*-terms such as radiology to represent their studies; the search terms used in this study would omit such studies. It is not expected however that there would be a set of terms that could be considered sufficient in retrieving research papers from a multidisciplinary area such as ehealth.

## Notes

### Competing Interest Statement

The authors have declared no competing interest.

